# Treatment of waste stabilization pond effluent using natural zeolite for irrigation potential

**DOI:** 10.1101/2021.10.25.465710

**Authors:** Kulyash Meiramkulova, Timoth Mkilima, Aliya Kydyrbekova, Elmira Bukenova, Abdilda Meirbekov, Gulnur Saspugayeva, Gulmira Adilbektegi, Kulzhan Beisembayeva, Gaukhar Tazhkenova

## Abstract

Direct utilization of treated effluent from natural treatment systems for irrigation can be challenging on sensitive plants due to high levels of salinity. Post-treatment of such an effluent prior to its applicability in irrigation can be of significant importance. In this study, the wastewater from a natural treatment plant was treated using a lab-scale filtration system with zeolite as a filter material. Three different column depths (0.5 m, 0.75 m, and 1 m) were used to investigate the effect of column depth on the treatment efficiency of the media. The suitability of the raw wastewater and the treated effluent from each column for irrigation purposes was investigated. The water quality parameters investigated were; electrical conductivity (EC), total dissolved solids (TDS), sodium (Na+), calcium (Ca2+), and magnesium (Mg2+). From the analysis results, it was observed that the column depth had a significant influence on the removal efficiency of the pollutants. Where the removal efficiency was observed to be increasing with the increase in the column depth. The highest removal efficiency (94.58%) was achieved from the combination of electrical conductivity and 1 m column depth, while the lowest removal efficiency (10.05%) was observed from the combination of calcium and 0.5 m column depth. The raw wastewater fell mostly into a “very high” hazard, which is class four (C4) based on electrical conductivity and class four (S4) based sodium adsorption ratio; making it unsuitable for irrigation purposes. However, the status improved after the treatment using different column depths.

## 1. Introduction

Effluent generated from natural wastewater treatment systems such as waste stabilization ponds is usually saline compared to freshwater due to high concentration levels of dissolved salts. Generally, natural treatment systems are those having little to no dependence on mechanical elements and chemicals to support wastewater purification. The systems mostly depend on plants and microorganisms such as bacteria to decompose and neutralize pollutants in wastewater [1–3].

Unfortunately, the linked high concentrations of salinity in the effluent could pose a significant threat to soil quality and plant growth at large upon its direct application in irrigation without being properly treated [4]. In general, soil salinization is regarded to be among the major threats to plant growth and affecting the agricultural sector in many parts of the world [5,6]. It is also important to note that, natural treatment systems are the most widely used technologies for wastewater treatment in the world [7–9], generating huge volumes of effluents every day. With the fact that freshwater is a vital and scarce resource [10], the reuse of effluents from natural treatment systems becomes an ideal solution for irrigation purposes.

However, studies have observed that crops grown on soils with high electrical conductivity (EC) can significantly reduce their yield [11]. Carbonates, chlorides, sulfates, and bicarbonates of sodium, potassium, magnesium, and calcium are among the salts that can be found in the effluent from waste stabilization ponds. Moreover, when the soil is more saline-sodic, the growth is affected by a combination of high alkalinity, high sodium (Na^+^), as well as high salt concentration [12]. Therefore, it is always necessary to differentiate between soil salinization and soil sodicity. However, salt tolerance among plants differs from one plant to another [13,14].

Generally, salt tolerance can be defined as the state at which a plant can grow and complete its life cycle on a substrate with high concentrations of levels of soluble salt [15]. Halophytes is the name given to plants that can withstand high concentrations levels of salt in the rhizosphere and grow well [16,17]. There are many crops with low salinity tolerance including rice (Oryza sativa L.) that has been observed to be highly susceptible to the rhizosphere salinity than other cereals [18]. From rice, it has been observed that high sensitivity mainly occurs at the vegetative and reproductive stages [19].

Among the challenges of salt, accumulation is the tendency of reducing the ability of the plants to uptake water and nutrients, resulting in osmotic or water-deficit stress. For sensitive plants, salt causes injury of the young photosynthetic leaves as well as accelerating their senescence. This is due to the fact that the Na+ cation accumulated in cell cyto-sol results in affecting the transpiration process of leaves [20]. To determine the suitability of water for use in irrigation, the sodium adsorption ratio (SAR) has been widely applicable as an indicator based on the concentrations of the main alkaline and earth alkaline cations present in the water [21]. Apart from EC and Na^+^, other parameters such as total dissolved solids (TDS), magnesium ion (Mg^2+^), and calcium ion (Ca^2+^) are also important to investigate the suitability of water for irrigation.

Therefore, to make the saline effluent from waste stabilization ponds reusable in irrigation, many treatment technologies have been introduced into the water industry. The technologies include the use of medium filtration and membrane filtration, cation exchange, electrodialysis, sorption, and electrochemical treatments. However, issues related to high capital costs especially due to energy consumption have hindered their application in wider regions [22].

This phenomenon brings significant importance to investigate the potential applicability of low-cost approaches for the treatment of biological treatment plant effluent with respect to irrigation purposes. The application of natural or synthetic zeolites as ion exchangers and adsorbents is regarded as among the relatively economical solutions for treating and reusing wastewater of high salinity [23]. Zeolites as a filter material provide one of the economical technologies and have been widely used in the field of water treatment as ion exchangers and adsorbents [24,25].

In this study, the potential applicability of zeolites on treating the effluent from a waste stabilization pond for irrigation purposes of low salinity tolerance plants is investigated. Three different column depths (0.5 m, 0.75 m, and 1 m) are used the investigate the influence of column depth on the treatment efficiency of zeolite. Then the effluent from each column is investigated for its potential applicability in irrigation, especially for sensitive plants.

## 2. Materials and Methods

### 2.1 Case study description, analytical methods, and wastewater characteristics

The wastewater samples used in this study were collected from the Vingunguti wastewater stabilization ponds in Dar es Salaam, Tanzania, approximately 7.2 km from the city Centre (latitude: 6°50’17.20”S, longitude: 39°14’8.62”E).

A number of water quality parameters were investigated in this study, including; EC, Na^+^, TDS, Mg^2+^, and Ca^2+^. The selection of the parameters is based on their significance in determining water suitability for irrigation purposes. In general, Na^+^ was measured using the Sodium-Ion Selective Electrode Method [26], while both Mg^2+^ and Ca^2+^ in water samples were measured using the Ethylenediaminetetraacetic Acid (EDTA) Method[27], with Na_2_EDTA 0.05, Acetylacetone, Tris(hydroxymethyl)-aminomethane (TRIS), Distilled Water and Electrolyte solution L300 as reagents. TDS and EC were determined using the TDS Meter Digital Water Tester (Lxuemlu, Shenzhen, China).

Table 1 presents a summary of the raw wastewater characteristics in terms of minimum (Min), maximum (Max) median, arithmetic mean (AM), and standard deviation (STD). A maximum EC concentration of 4224 µS/cm was observed from the raw wastewater, with an average concentration of 2478.1 µS/cm.

**Table 1.** Raw wastewater characteristics

| Parameter | Min | Max | Median | AM | STD |
| --- | --- | --- | --- | --- | --- |
| EC | 1001 | 4224 | 2251 | 2478.10 | 731.975 |
| TDS | 996 | 2284 | 1602 | 1622.20 | 332.429 |
| Na<sup>+</sup> | 115.5 | 145.7 | 133.8 | 131.93 | 9.226 |
| Mg<sup>2+</sup> | 8.7 | 22.5 | 15.1 | 14.61 | 4.095 |
| Ca<sup>2+</sup> | 6.2 | 14.3 | 8.85 | 9.30 | 2.344 |

EC in µS/cm, all other parameters in mg/L

### 2.2 Experimental setup

Three different fixed-bed columns with 0.5 m, 0.75 m, and 1 m depths were used to investigate the influence of column depth in the treatment efficiency of natural zeolite (Figure 1). The column containers are of Polyvinyl chloride (PVC) material with approximately 5.08 cm in diameter. All three columns were packed with natural zeolite adsorbents (clinoptilolite) composed of a microporous arrangement of silica and alumina tetrahedra with an average particle size of 1.5 mm (FM Stock and Supplies, Kenmare, Gauteng, South Africa).

**Figure 1.**
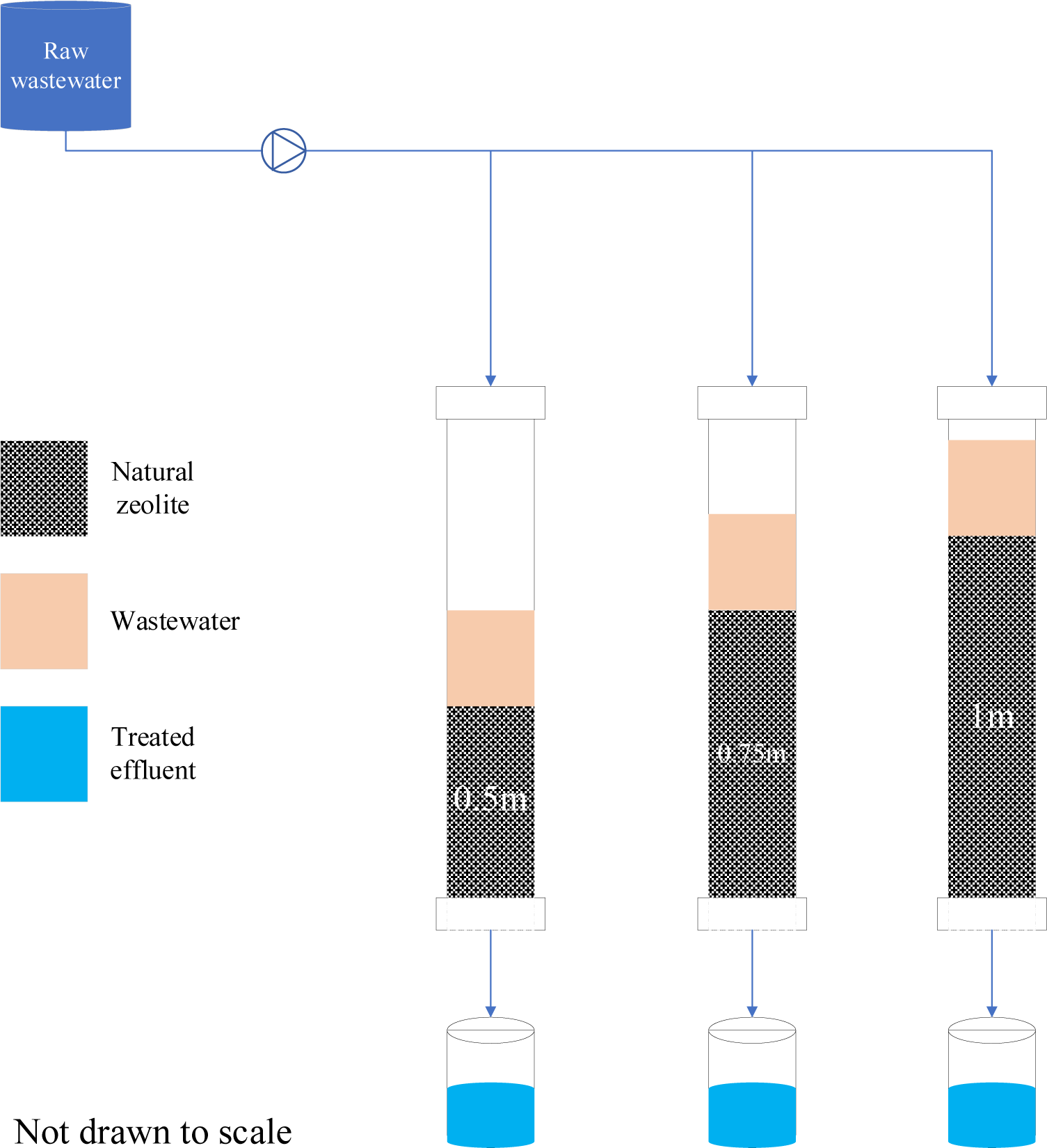
Experimental setup

To allow equal distribution of flow in the columns, the top surfaces of the columns were covered by perforated plates with evenly distributed holes. 100 L storage drum was used to feed the columns at a controlled rate of 0.0035 L/s. To maintain all the solids in suspension, the wastewater was slowly and continuously stirred. The Wet-packing approach of the porous medium was used with the purpose of minimizing layering and air entrapment inside the filing. All three columns were mounted vertically, and glass wool was used at the bottom of the column acting as supporting material of the adsorbent bed. After packing the column, deionized water was passed through the column for some time, followed by the introduction of the feed water. The filtrate samples were collected at a regular time interval. All the experiments were carried out at room temperature (20 to 25 ^°^C).

#### Statistical methods

Correlation analysis was among the approaches used to analyze results from the experiments. Correlation matrices were developed to evaluate the strength of the relationship among the studied parameters. A high correlation indicates that two or more variables have a strong relationship with each other. While a weak correlation provides an indication that the variables are hardly related. Table 2 provides a summary of the interpretation of the correlation indices used in this study.

**Table 2.** Interpretation of the correlation coefficients

| Range of correlation coefficient | Strength of relationship |
| --- | --- |
| 0-0.29 | Weak |
| 0.3-0.49 | Moderate |
| 0.5-0.69 | Strong |
| 0.7-1 | Very strong |

Apart from the correlation matrices, box and whisker plots were used to evaluate data distributions among the water quality parameters. The evaluation is based on the distribution of numerical data and skewness through data quartiles (percentiles) and averages. In general, box plots show the five-number summary of a set of data: the minimum score, first (lower) quartile, median, third (upper) quartile, and maximum score. In this study; EC, Na, Mg, Ca, and TDS were analyzed using box and whisker plots.

The salinity hazard zones based on electrical conductivity (EC) were classified into four classes; class one (C1) class two (C2), class three (C3), and class four (C4), ranging from low salinity (C1) to very high salinity (C4). Table3 provides a summary of the salinity hazard zones based on EC with their interpretations in terms of usability.

Table 4 provides a summary of the salinity hazard zones based on Sodium adsorption ratio (SAR) with their interpretations in terms of usability. The salinity hazard zones based on SAR were also classified into four classes and used in the Wilcox diagrams; class one (S1) class two (S2), class three (S3), and class four (S4) ranging from low sodium hazard (S1) to very high sodium hazard (S4).

**Table 3.** Salinity Hazard Zones: based on electrical conductivity

| Water class | EC (μS/cm) | Definition |
| --- | --- | --- |
| C1 - Low salinity | 0-250 | Water can be used safely |
| C2 - Medium salinity | 250-750 | Water can be used with moderate leaching |
| C3 - High salinity | 750-2250 | Water can be used for irrigation purposes with some management practices |
| C4 - Very high | 2250-5000 | Water cannot be used for irrigation purposes |

**Table 4.** Sodium Hazard Zones: based on Sodium Adsorption Ratio lines

| Water class | SAR | Definition |
| --- | --- | --- |
| S1 low sodium hazard | 0-10 | Little or no hazard |
| S2 medium sodium hazard | (10-18) | Appreciable hazard but can be used with appropriate management |
| S3 High sodium hazard | 18-26 | Unsatisfactory for most of the crops |
| S4 Very high sodium hazard | > 26 | Unsatisfactory for most of the crops |

SAR can be defined as an index used to define the effect of sodium concentration in a sample in relation to calcium and magnesium. More specifically, the SAR index is achieved by diving the square root of 1/2 of the calcium plus magnesium concentrations. Equation 1 provides a summary of the formula used in the computations of SAR [28].

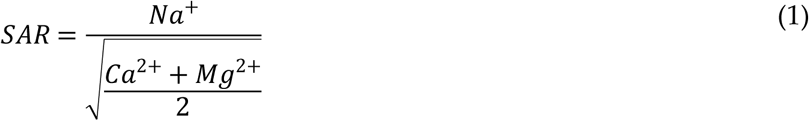

Furthermore, Wilcox diagrams were plotted from the raw wastewater, effluent from 0.5 m, 0.75 m, and 1 m zeolite columns. By definition, the Wilcox plot is a semi-log scatter plot of the “sodium hazard” (SAR) on the Y-axis versus the “salinity hazard” (EC) on the X-axis. It has to be noted that, the EC is plotted by default in a log scale. The treated effluent suitability for irrigation mainly depends on the concentration of total salinity and sodium related to other ions [29]. Therefore, the diagrams were used to evaluate the risk levels in the raw wastewater and the treated effluent from the three columns.

## 3. Results and discussion

The analysis of the water samples before and after the treatment was successfully executed. In the raw wastewater samples, the average EC concentration was 2478.1 µS/cm, while that of TDS was 1622.20 mg/L. The EC concentration in the raw wastewater falls in class four (C4) based on the salinity hazard zones, with an indication that the effluent from waste stabilization ponds cannot be used directly for irrigation purposes. Average concentrations of 131.9 mg/L, 14.6 mg/L and 9.3 mg/L were recorded from Na^+^, Mg^2+^, and Ca^2+^, respectively.

From Figure 2, it can be observed that from the EC boxplot, the median line is closer to the middle, indicating that the EC data distribution is symmetric or normal. From the Na^+^ boxplot, it can be observed that the median line closer to the upper quartile with an indication that the distribution of Na^+^ data in the raw wastewater is considered to be “negatively skewed”. This means the Na^+^ data constituted a higher frequency of low concentration values than the high concentration values.

**Figure 2.**
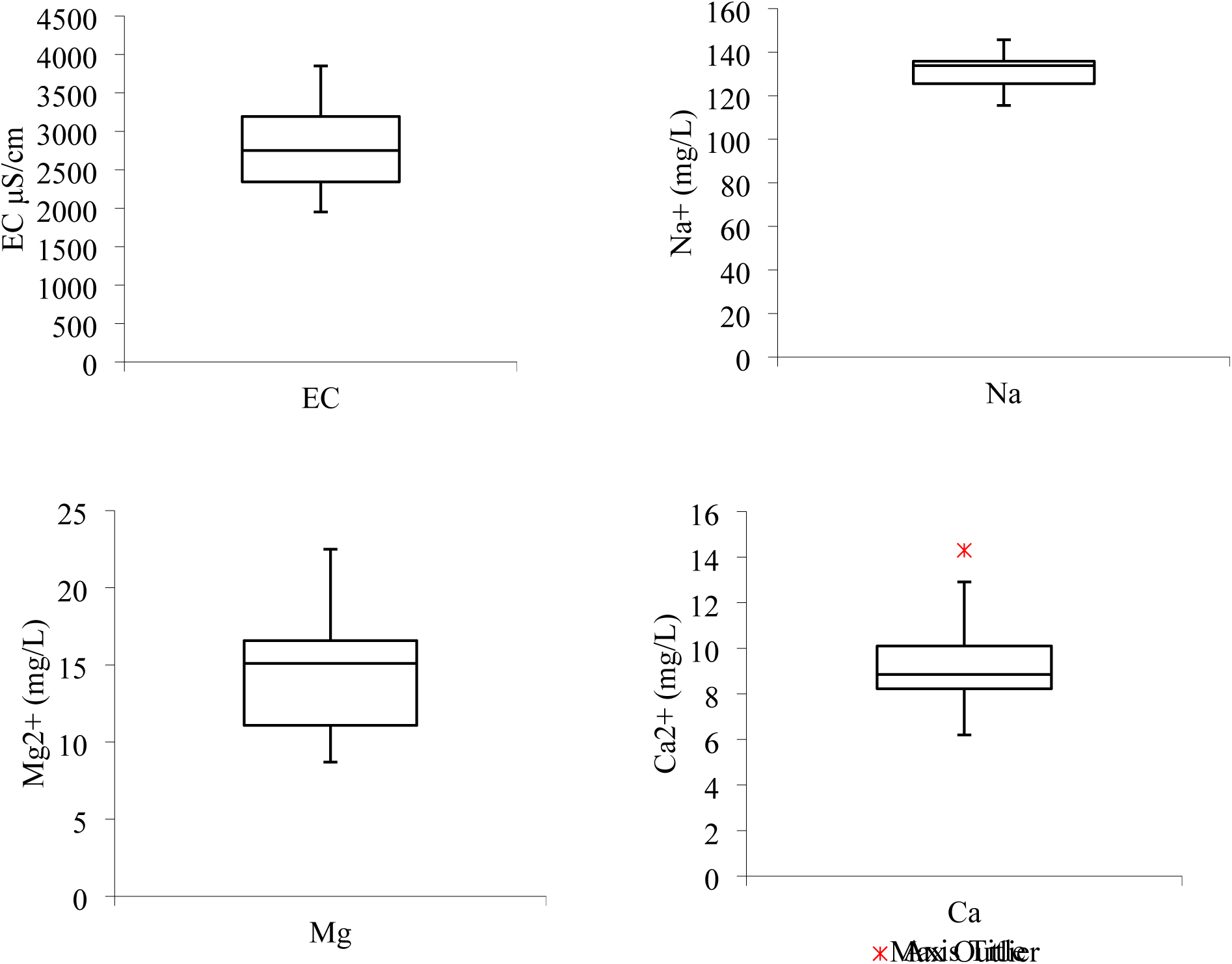
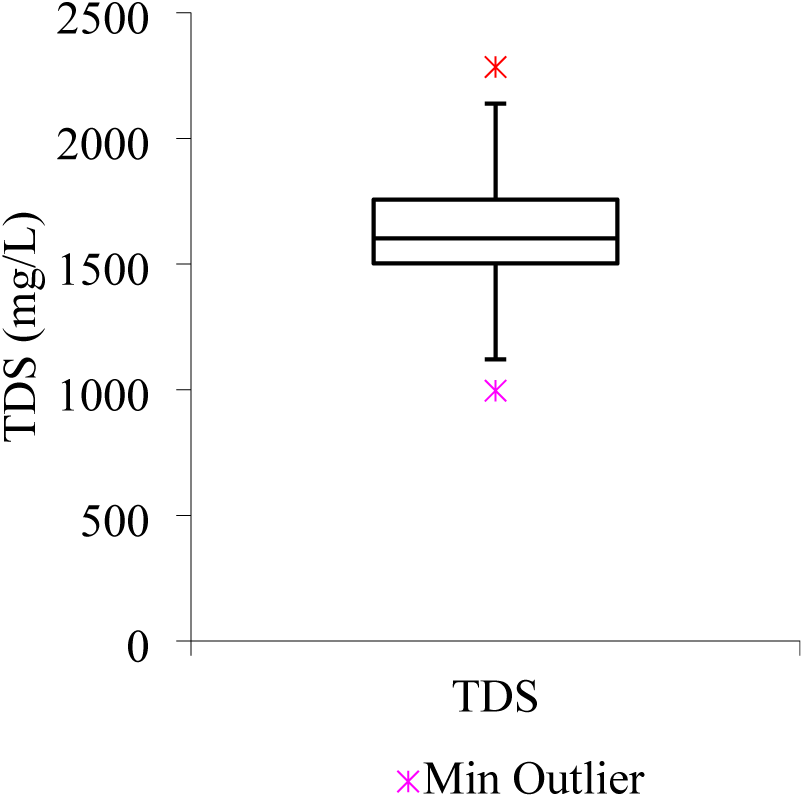
Boxplots from raw wastewater (a) EC (b) Na^+^ (c) Mg^2+^ (d) Ca^2+^ (e) TDS

As observed from the Na^+^ boxplot, a similar case applies to Mg^2+^ concentration data distribution from the raw wastewater. While, from the Ca^2+^ boxplot, the median line is observed to be closer to the lower quartile meaning that the water quality data constitute a higher frequency of more high concentration values than the low concentration values also known as “positive skewness”. Similarly, from the TDS boxplot, the median line is observed to be closer to the lower quartile meaning that the water quality data constitute a higher frequency of high concentration values than the low concentration values (“positively skewed”).

A correlation matrix for the studied water quality parameters in raw wastewater was developed to evaluate the strength of the relationship among them. From Table 5, it can be observed that the general correlation among the parameters ranging from a strong to a very strong relationship. The highest correlation index of 0.966002 was achieved between Na and EC, followed by 0.945631 between TDS and Na. Also, a very strong correlation can be observed between TDS and EC with a correlation index of 0.944682. The lowest correlation index can be observed between the Mg^2+^ and Na^+^ with a correlation index of 0.598734. However, the index between Mg^2+^ and Na^+^ falls under a strong relationship.

**Table 5.** Correlation matrix from raw wastewater

|  | EC | Na<sup>+</sup> | Mg<sup>2+</sup> | Ca<sup>2+</sup> | TDS |
| --- | --- | --- | --- | --- | --- |
| EC | 1 |  |  |  |  |
| Na<sup>+</sup> | 0.966002 | 1 |  |  |  |
| Mg<sup>2+</sup> | 0.650596 | 0.598734 | 1 |  |  |
| Ca<sup>2+</sup> | 0.837318 | 0.766448 | 0.725882 | 1 |  |
| TDS | 0.944682 | 0.945631 | 0.611418 | 0.865351 | 1 |

In the literature, other studies have also observed a very strong relationship between TDS and EC to the point of recommending the EC to estimate TDS based on the linear relationship as shown in Equation 2 [30,31]. With the fact that TDS measurement is considered to be a time-consuming process, simplicity is often estimated from EC assuming TDS are predominantly ionic species of low enough concentration to produce a linear TDS-EC relationship [32].

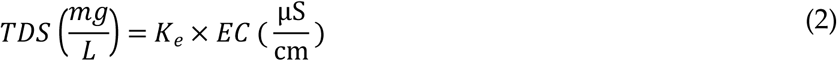

Where; K_e_ is a proportionality constant ranging from 0.54 to 1.1.

According to Thirumalini and Joseph [33], that investigated the correlation between EC and TDS in natural waters, it was observed that the correlation index between TDS and EC was 0.63 for samples taken from pollution-free residential areas as well as ranging from 0.59 to 0.93 for samples taken from the textile industrial belt. Therefore, the general strong correlation between TDS and EC observed in the literature agrees with the results obtained from this study.

From Tables 6 to 8, it can be observed that, when the depth was increased from 0.75 m to 1 m, there was slight difference in terms of EC and TDS removal in the wastewater. The average EC concentration from the 0.75 m column depth was 487.85 µS/cm, while from 1 m column depth, the average EC concentration was 378.51 µS/cm. Also, the average TDS concentration from the 0.75 m column depth was 134.36 µS/cm, while that of 1 m column depth was 130.163 µS/cm. The phenomenon suggests that, the treatment approach has a gradual removal efficiency as the column depth increases from 0.75 m to 1 m.

**Table 6.** Water quality characteristics from 0.5 m column depth effluent

| Parameter | Min | Max | Median | Mean | STD |
| --- | --- | --- | --- | --- | --- |
| EC | 901 | 2403 | 1202 | 1352.898 | 432.395 |
| TDS | 980 | 1282 | 1101 | 1107.3 | 102.363 |
| Na | 86.6 | 127.6 | 105.65 | 106.38 | 11.295 |
| Mg | 6.4 | 10.6 | 9 | 8.87 | 1.416 |
| Ca | 4.4 | 10.3 | 8.95 | 8.365 | 1.918 |

**Table 7.** Water quality characteristics from 0.75 m column depth effluent

| Parameter | Min | Max | Median | Mean | STD |
| --- | --- | --- | --- | --- | --- |
| EC | 310 | 662 | 501 | 487.8531 | 98.452 |
| TDS | 88 | 239.5 | 123 | 134.36 | 42.892 |
| Na | 52.9 | 72.8 | 63.75 | 64.07 | 7.182 |
| Mg | 2.06 | 10.4 | 6.45 | 6.51 | 2.383 |
| Ca | 4.5 | 11.6 | 7.6 | 7.81 | 2.463 |

**Table 8.**
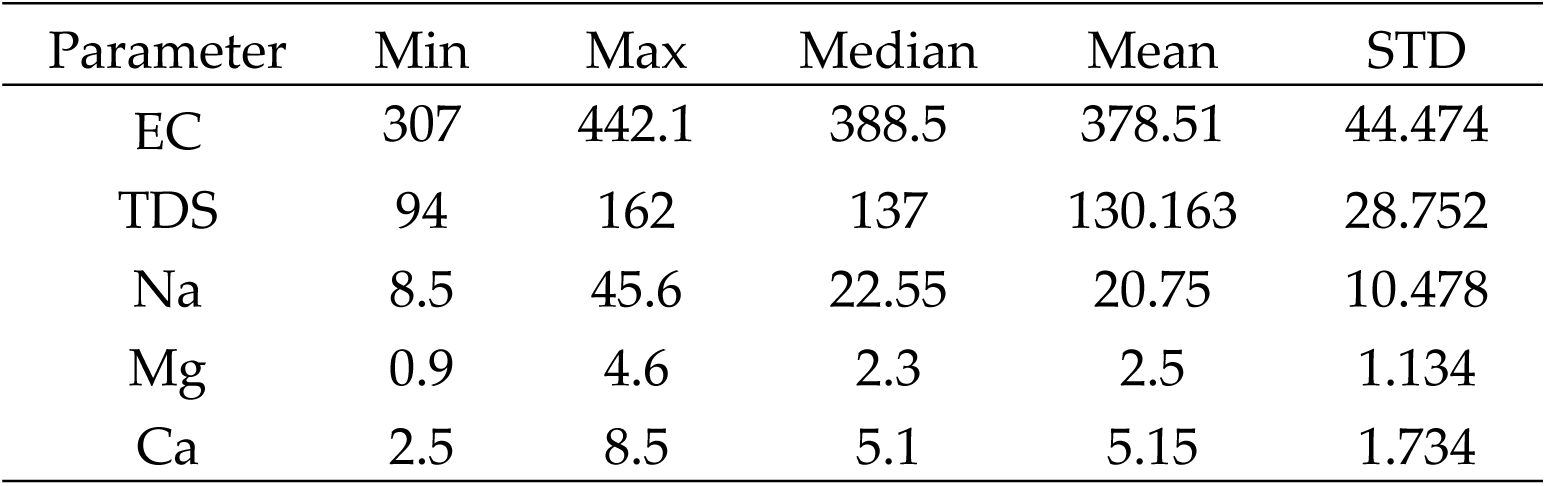
Water quality characteristics from 1 m column depth effluent

| Parameter | Min | Max | Median | Mean | STD |
| --- | --- | --- | --- | --- | --- |
| EC | 307 | 442.1 | 388.5 | 378.51 | 44.474 |
| TDS | 94 | 162 | 137 | 130.163 | 28.752 |
| Na | 8.5 | 45.6 | 22.55 | 20.75 | 10.478 |
| Mg | 0.9 | 4.6 | 2.3 | 2.5 | 1.134 |
| Ca | 2.5 | 8.5 | 5.1 | 5.15 | 1.734 |

From Figure 3, the EC, Na^+^ and TDS boxplots, the median line closer to the upper quartile with an indication that the distribution of EC, Na^+,^ and TDS data in the treated effluent using the 1 m column of zeolite is considered to be “negatively skewed”. While that of Mg^2+^ is observed to be closer to the middle, indicating that the Mg^2+^ data distribution was symmetric or normal. From the Ca^2+^ boxplot, the median line is seen to be closer to the upper quartile with an indication that the distribution of Ca^2+^ data in the treated effluent using the 1 m column of zeolite is considered to be “negatively skewed”. This means the Ca^2+^ data constituted a higher frequency of low concentration values than the high concentration values.

**Figure 3.**
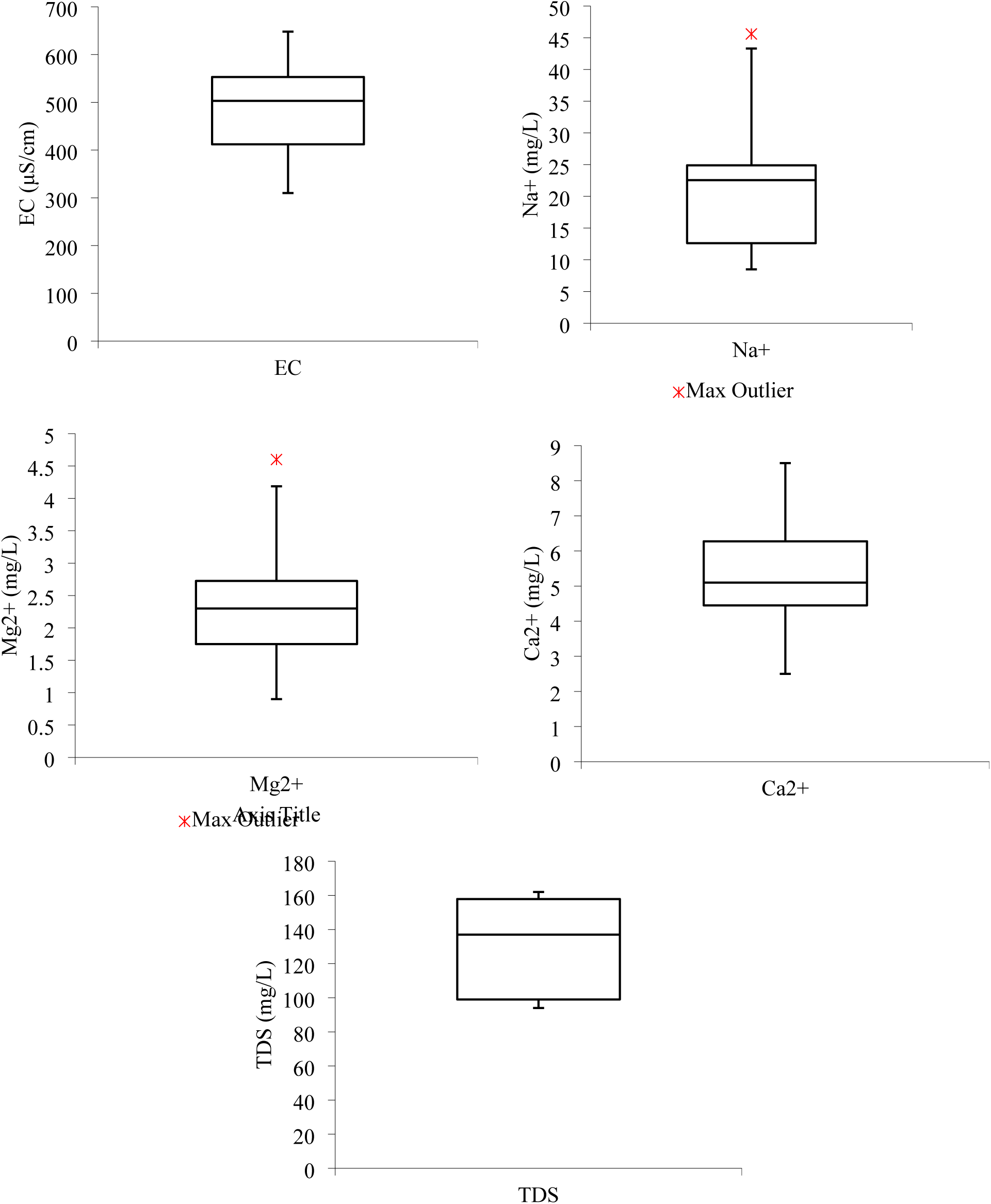
Boxplots from 1 m column effluent (a) EC (b) Na^+^ (c) Mg^2+^ (d) Ca^2+^ (e) TDS

The removal efficiency of the studied parameters was observed to be significantly affected by the column depths; the more the column depth, the higher the removal efficiency. The highest removal efficiency (94.58%) was achieved from the combination of EC and 1 m column depth. This was followed by a removal efficiency of 91.98% from the combination of TDS and 1 m column depth. The lowest removal efficiency can be observed from the combination of Ca^2+^ and 0.5 m column depth. In the literature, natural zeolite has also been observed to be highly efficient in terms of TDS removal. According to [34], which investigated the treatability of brackish groundwater by zeolite filtration in Sumur Tua Wonocolo, Kedewan, Bojonegoro, East Java, a TDS removal efficiency of up to 84% was achieved.

From Figure 5, it can be observed that the raw wastewater falls under high (C3) to very high (C4) hazards based on the EC. Approximately 68% of the values fall under the very high hazard while 32% fall under the high hazard category. While, based on the SAR, the raw wastewater falls from low (S1) to very high (S4), with the majority of the values falling under medium (S2) and high (S3). The general phenomenon suggests that raw wastewater is not recommended for irrigation purposes especially for low-salt-tolerance plants based on EC and unsatisfactory for most of the crops based on SAR.

**Figure 4.**
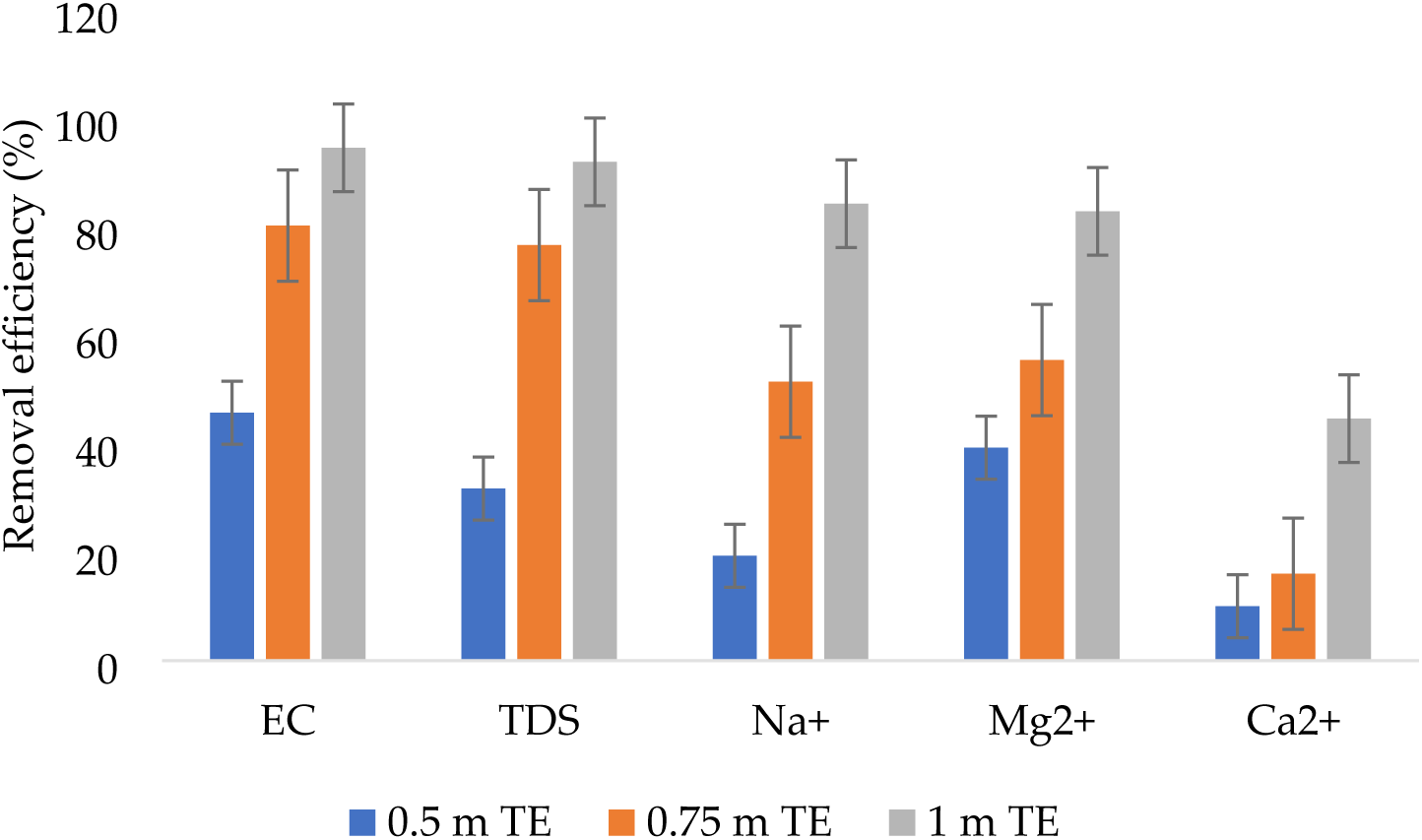
Removal efficiencies

**Figure 5.**
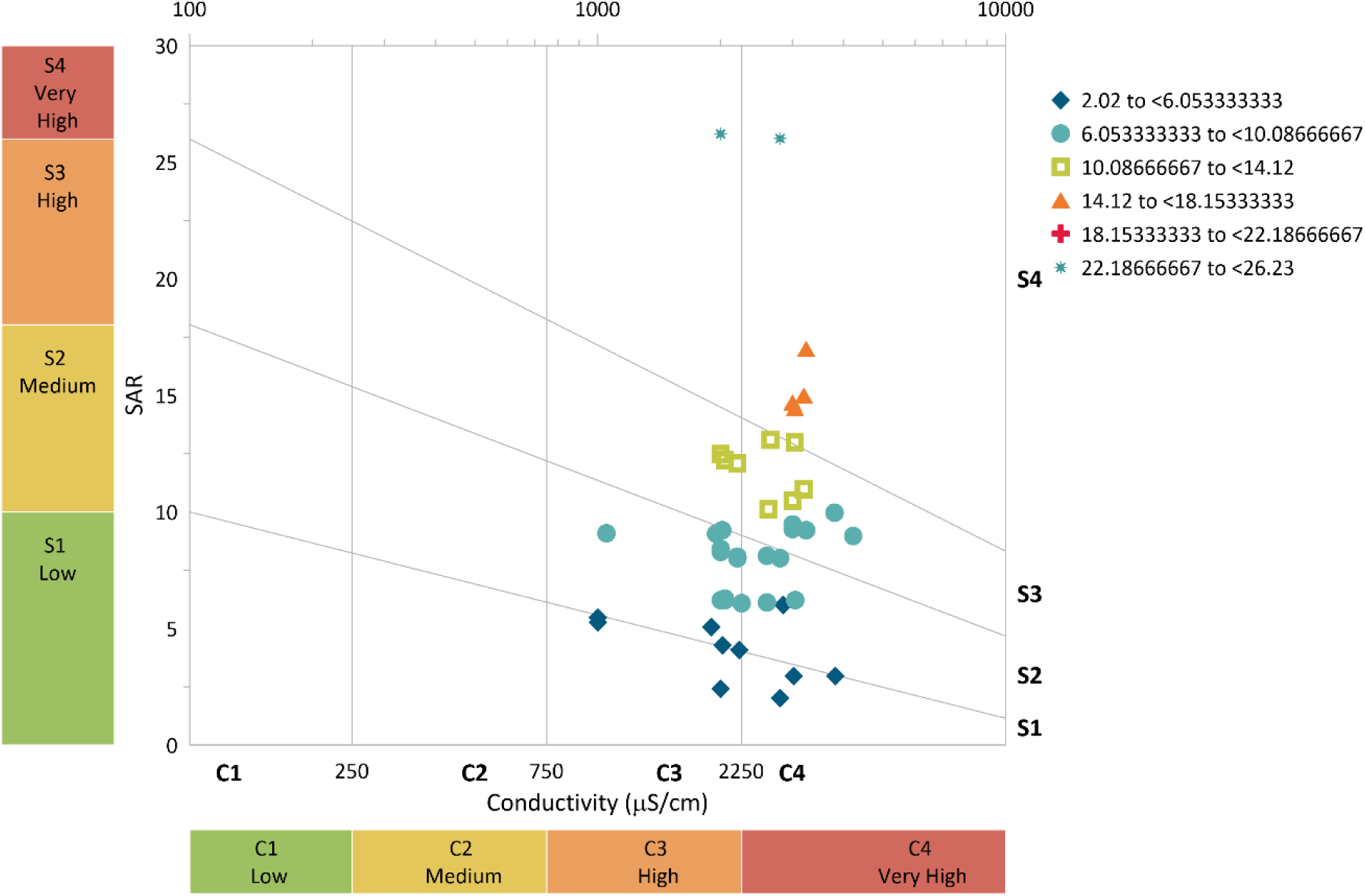
Salinity hazard from raw wastewater

From Figure 6, it can be observed that the treated effluent from 0.5 m column of zeolite falls under high (C3) to very high (C4) hazard based on the EC. Approximately 97% of the values fall under the high hazard while 3% fall under the very high hazard category. While, based on the SAR, the treated effluent from 0.5 m zeolite column falls under low (S1) to high (S4), with the majority of the values falling within medium (S2) hazard. The general phenomenon suggests that the treated effluent from the 0.5 m column is under appreciable hazard but can be used with appropriate management based on SAR and can be used for irrigation purposes with some management practices based on EC.

**Figure 6.**
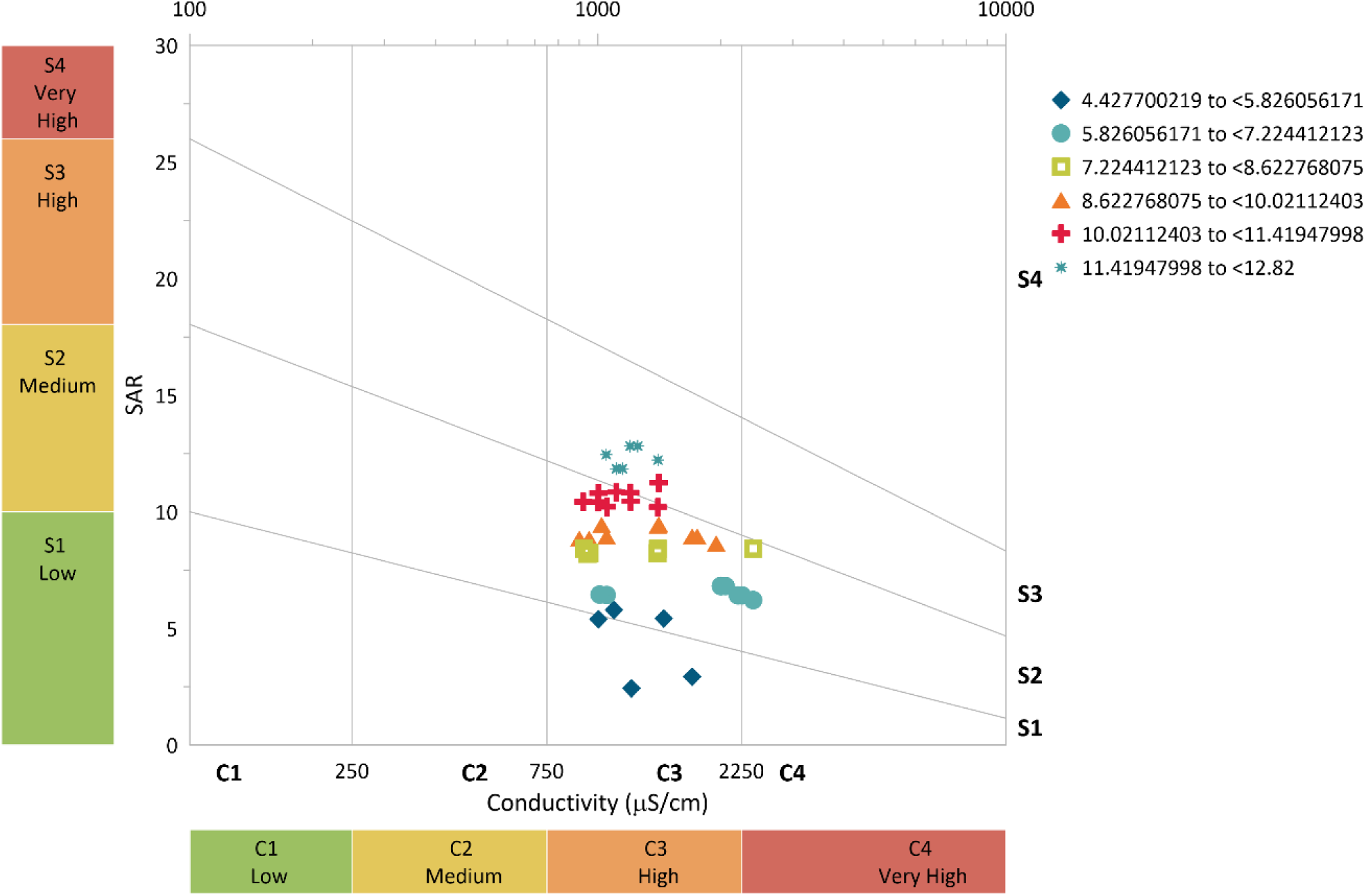
Salinity hazard from 0.5 m column effluent

From Figure 7, it can be observed that the treated effluent from the 0.75 m zeolite column falls under medium (C2) hazard based on the EC. Almost 100% of the values fall under medium hazard. While, based on the SAR, the treated effluent from 0.75 m falls under low (S1), with very few values getting close to medium (S2). The general phenomenon suggests that the treated effluent from a 0.75 m column can be used for crop irrigation purposes with little or no hazard based on SAR and can be used with moderate leaching based on EC.

**Figure 7.**
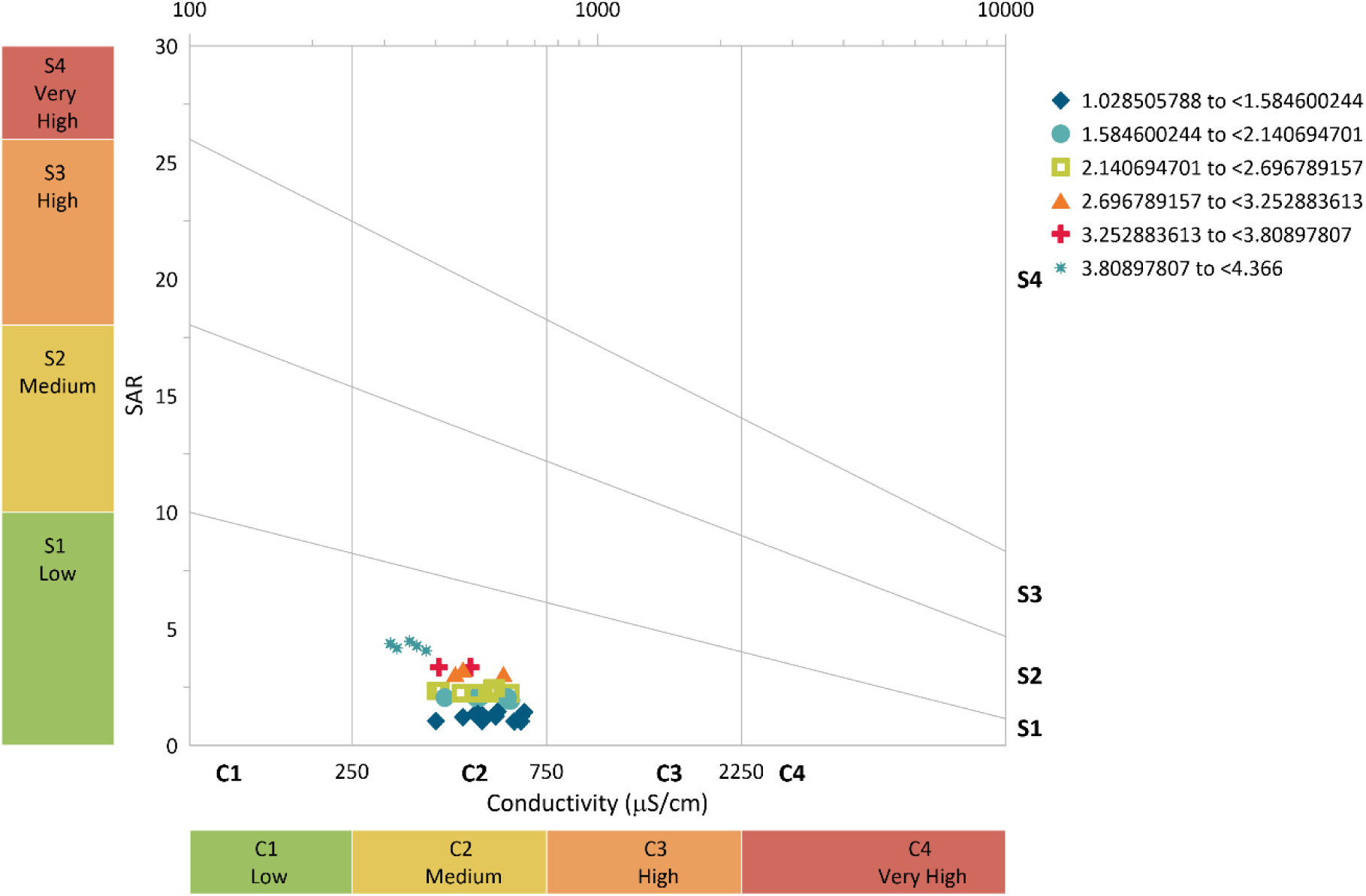
Salinity hazard from 0.75 m column effluent

From Figure 8, it can be observed that the treated effluent from the 1 m zeolite column falls under a low (C1) hazard based on the EC. While, based on the SAR, the treated effluent from 1 m column under low (S1), with very little values getting close to medium (S2). The general phenomenon suggests that the treated effluent from the 1 m column can be used for crop irrigation purposes with little or no hazard based on SAR and can be used safely based on EC.

**Figure 8.**
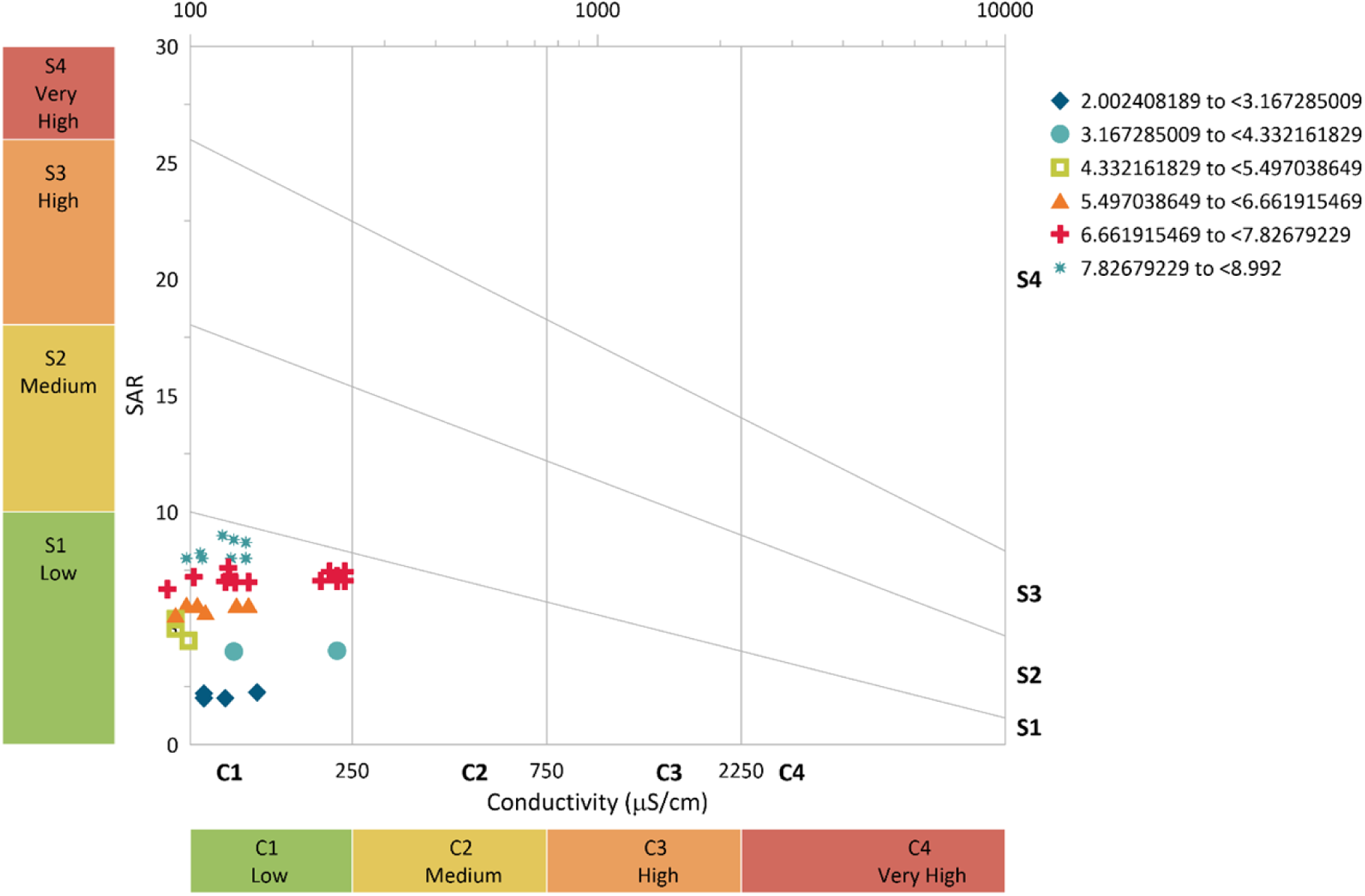
Salinity hazard from 1 m column effluent

## 4. Conclusions

In this study, the potential applicability of zeolites on treating the effluent from a waste stabilization pond for irrigation purposes has been investigated with three different column depths. A correlation among the studied parameters was observed with the highest correlation index of 0.966002 achieved between Na^+^ and EC, followed by 0.945631 between TDS and Na^+^. Also, the results showed that the pollutants removal efficiency increased with the increase in column depth. Among the studied parameters, the highest removal efficiency (94.58%) was achieved from the combination of EC and 1 m column depth, while the lowest removal efficiency (10.05%) was observed from the combination of Ca^2+^ and 0.5 m column depth. From the hazard analysis, the raw wastewater generally fell into the “very high” hazard class based on both EC and SAR. In that matter, the raw wastewater has to be treated further prior to its application for irrigation purposes. However, the status improved after the treatment using different column depths. In that matter, the results from this study revealed further that, it is always important to investigate the quality of effluent from natural treatment systems before subjecting it to any sort of irrigation. Moreover, the use of zeolite can provide one of the efficient approaches to improve the effluent and make it suitable for irrigation.

## Author Contributions

Conceptualization, T.M; methodology; software, T.M. and K.M., validation, K.M., G.A., and G.S.; formal analysis, E.B.and A.K.; investigation, T.M.; resources, K.M.; data curation, K.M., A.M., G.T., K.B., T.M., and To.M.; writing—original draft preparation, T.M., writing—review and editing T.M. and K.M.; visualization.; supervision, K.M.; project administration K.M., All authors have read and agreed to the published version of the manuscript.

## Data Availability Statement

Not applicable

